# DatumKB: A Database of Biological Experimental Results

**DOI:** 10.1101/2021.06.25.449966

**Authors:** Merrill Knapp, Tim McCarthy, Carolyn Talcott

## Abstract

1.

DatumKB is a freely accessible database of experimental results involving the function and regulation of human proteins in cultured cells. The results are manually curated from biological research literature using a shorthand language and stored as datums. Datums were originally designed to be used as evidence for rules in a Pathway Logic model of intracellular signal transduction (STM8, http://pl.csl.sri.com/online.html) (1). They are independent units that can be understood by biologists, traced back to their source, and have enough structure to be interrogated computationally. The information is expressed using a controlled vocabulary with links to well known databases such as HUGO Gene Nomenclature Committee (HGNC), UniProt, PubChem, and Cellosaurus. DatumKB can be searched using a query interface and the results can be downloaded in the original datum format, a simplified text format, or a JSON file. Links to detailed documentation of datum structure and a tutorial for the search engine are provided.

**Database URL:** https://datum.csl.sri.com

## 2. INTRODUCTION

One of the unaccomplished tasks of today is to implement a system that can help scientists collect, curate, annotate and validate experimental results found in published scientific literature.

There are many databases that collect the conclusions of a paper as annotations and enter them into an appropriate field of a database. The UniProt database (https://www.uniprot.org) uses this method to extract protein function, interactions, and subcellular locations and store them as free text (2). The Gene Ontology (http://geneontology.org) (3) uses a controlled vocabulary and an ontology to further categorize these properties. However, both of these approaches depend on the curator to translate experimental results into annotations. To learn how an attribute was determined experimentally, a user needs to go to the original reference and find the relevant experiments. In some cases, a more detailed representation of the data is needed.

What is an experiment? To a cell biologist studying intracellular signal transduction, an experiment begins with seeding cultured cells and setting up a starting state by making the cells quiescent, differentiating them into a certain cell type, or just seeding them so they will be exponentially growing at the time of treatment. After the cells are treated, a change in the state is determined by comparing the treated cells with untreated cells. Knowing the environment or and starting state of any experiment is crucial to the interpretation of the results; however, many repositories of high throughput data do not provide a structured way to store that information with the data. Consider the MIAME and MINSEQ guidelines for submission of data to The Gene Expression Omnibus database (https://www.ncbi.nlm.nih.gov/gds) at the US National Center for Biotechnology Information (NCBI) (4). They encourage the submitter to provide “Essential laboratory and data processing protocols” but are unclear about what these elements are. In contrast, the Encyclopedia of DNA Elements (ENCODE) (https://www.encodeproject.org) Data Coordinating Center (DCC) (5, 6) has created a system for organizing submitted data based on experiments. An experiment contains replicates of only one assay, one kind of biosample, and one target. This facilitates the searching for datasets based on how the data was obtained rather than on free-text descriptions of the results.

For low-throughput data reported in the published literature, specialized databases collect experimental results as evidence for phenomena such as protein-protein interactions or modification sites. BioGrid (https://thebiogrid.org) (7) uses a library of 20 physical experimental systems that are assigned to each protein-protein interaction. The IntAct Molecular Interaction Database (https://www.ebi.ac.uk/intact/) (8) uses the HUPO-PSI Controlled Vocabulary for interaction detection. These databases provide information about existing interactions, but they do not cover cases where an interaction is changed in response to a stimulus or mention interactions that could not be detected.

When we decided to make a model of intracellular signal transduction based on experimental evidence rather than curated assertions we needed a structure that describes the results of all the experiments in publications that are relevant to our model. Because the model rules are created manually from the experimental results, the system needs to store the evidence in a human-readable format and in a structure where each result can be visualized as a single unit. The units have to be written in a way that can be parsed by computers to provide a system for sorting, storage and retrieval. We also needed a system to map the experimental results back to the original published experiment so results can be reexamined when new information becomes available. The datum became our solution to these challenges.

## 3. DATUM STRUCTURE

A datum is a structured, computer readable summary of an experimental finding representing a single biological assay. The information in a datum includes the following elements:

- the Subject (a protein, gene, or cellular phenotype)
- the Assay (what was measured and how it was detected)
- the Change (increase, decrease, no change)
- the Stimulus (addition of a drug, peptide, or stress, and for how long)
- the Environment (cells and culture conditions used at the start of a treatment)
- the Source (a PubMed ID and the number of the figure or table containing the experimental result)

Datums have a syntax that allows them to be parsed by computer and translated into JSON for storage and retrieval (Figure 1). The **subject** can be a protein or a gene. When the subject is a protein, it must have a handle attribute that describes the way it was detected. The most common handles are “Ab” for an antibody against the endogenous protein or “tAb” for an antibody against the tag on an expressed protein. If the subject is a protein that has been immunoprecipitated the attribute “IP” is added. When a cellular phenotype is being detected, the subject is “NS” (for no subject) and the phenotype is described in the assay element. The different subject feature are illustrated in datums #1-3 below. The subject of datum #1 is exogenous S6k1 immunoprecipitated using an antibody against a tag (indicated by the prefix x) added to the expressed protein. The datum reports that the phosphorylation of S6k1 on T412 is increased in response to addition of Tnf to MCF cells for 30 minutes, The subject of datum #2 is the endogenous protein Myc as detected by an antibody. The datum reports an increase in endogenous Myc in response to addition of Egf to HCT116 cells. Datum #3 reports an experiment measuring a phenotype — an increase in apoptosis over time after treatment with Cisplatin.

**Figure 1:**
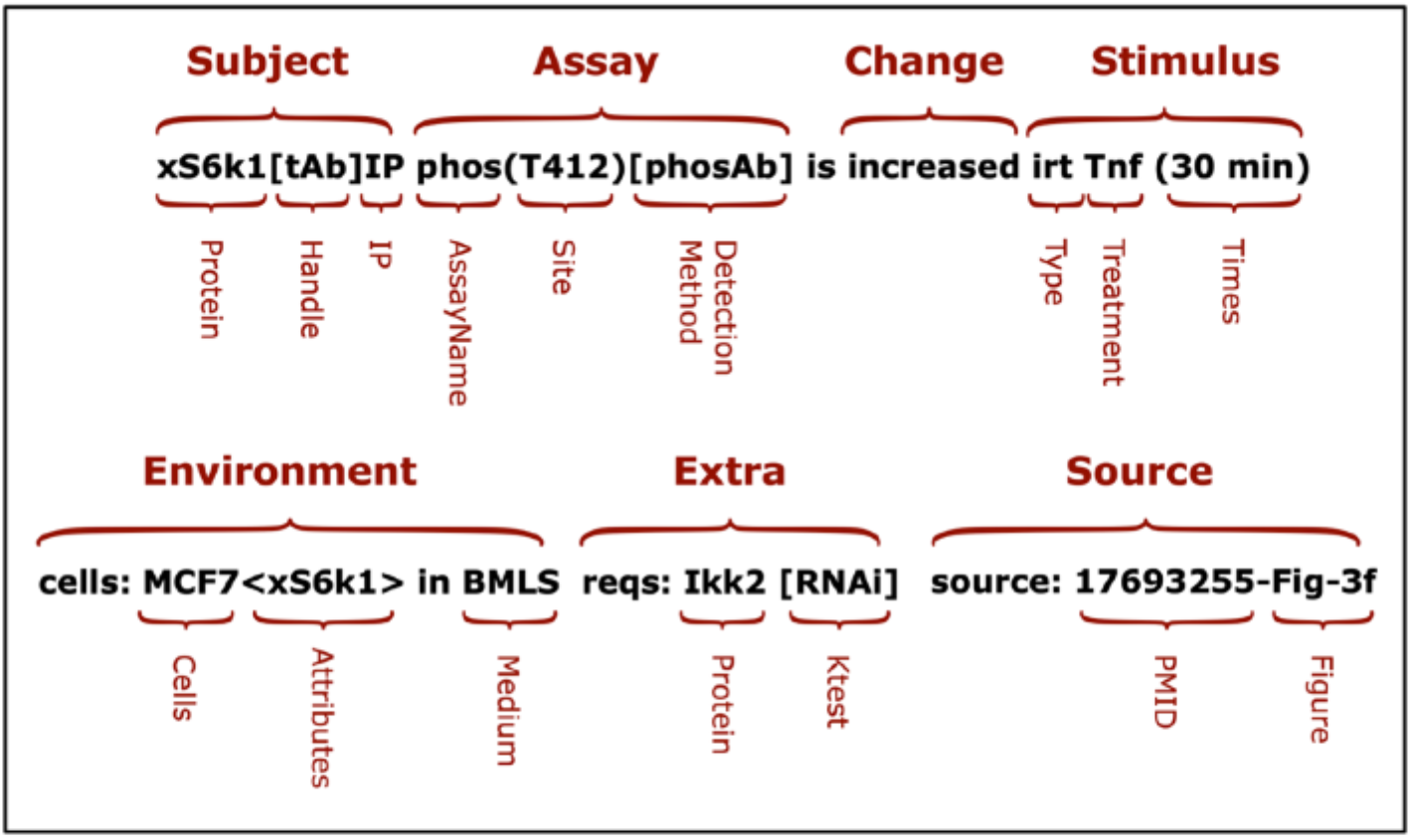
The Parts of a Datum

Datum #1:

~~~
 *** xS6k1[tAb]IP phos(T412)[phosAb] is increased irt Tnf (30 min)
   *** cells: MCF7<xS6k1> in BMLS
   *** reqs: Ikk2 [RNAi]
   *** source: 17693255-Fig-3f
~~~

**Figure 2:**
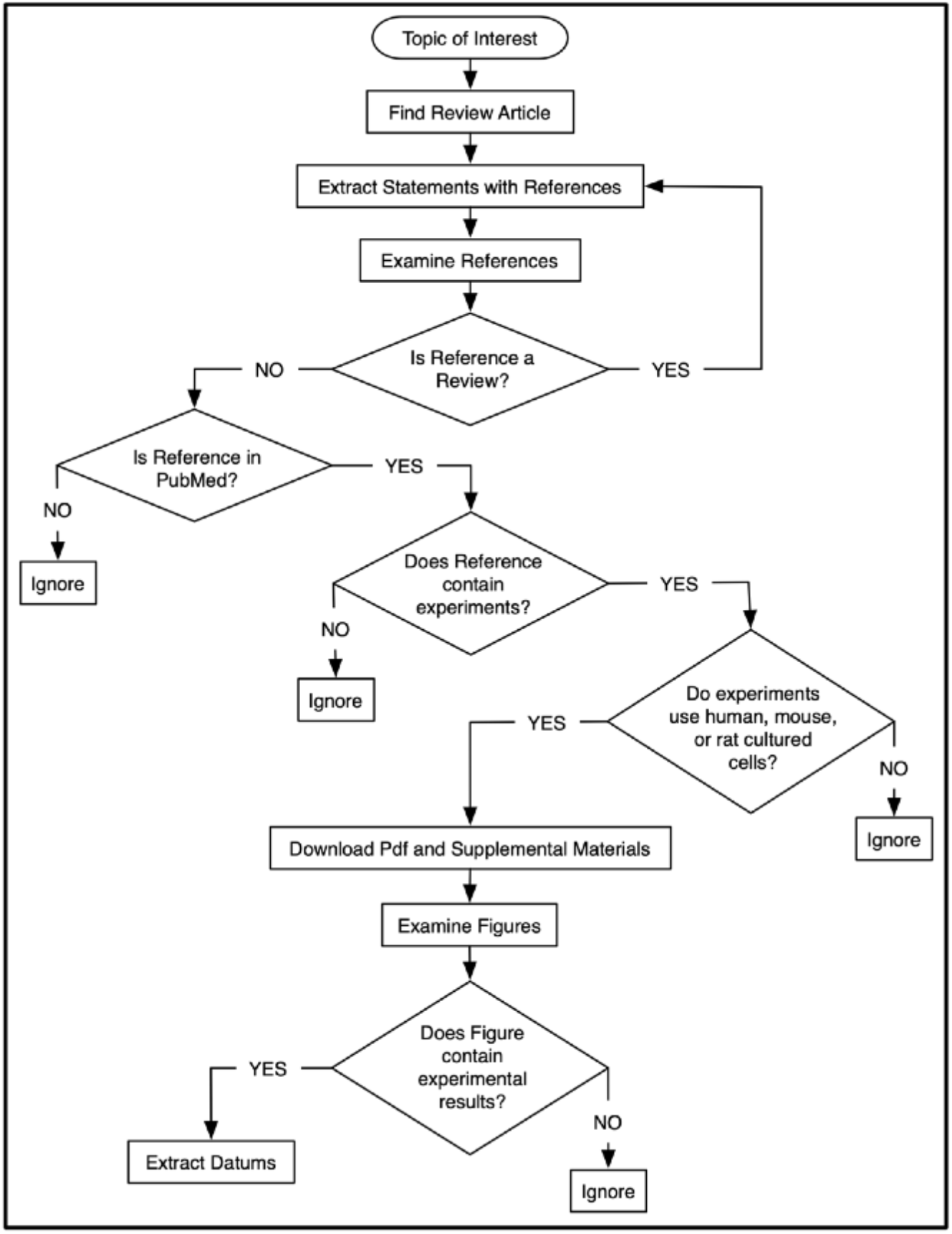
A workflow for choosing experiments to curate

**Figure 3:**
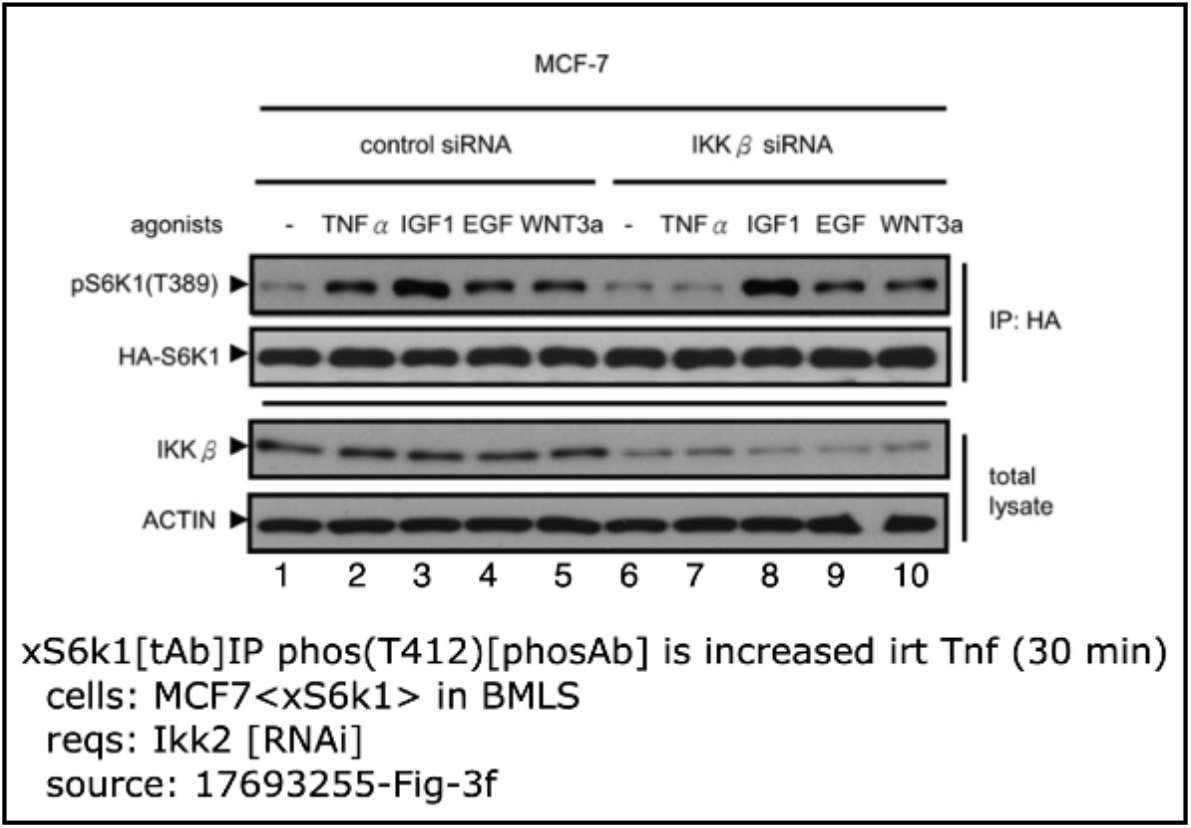
Example of a datum with the figure from which it was derived. Image reprinted from Cell 3:440-55 (2007) Lee,D.F., Kuo,H.P., Chen,C.T. et al. IKK beta suppression of TSC1 links inflammation and tumor angiogenesis via the mTOR pathway with permission from Elsevier.

Datum #2:

~~~
 *** Myc[Ab] prot-exp[WB] is increased irt Egf (3 hr)
   *** cells: HCT116 in BMLS
   *** reqs: Ctnnb1 [RNAi]
   *** source: 18356165-Fig-3c
~~~

Datum #3:

~~~
 *** NS Apoptosis[AnnexinV] is increased irt Cisplatin (times)
   *** cells: HCT116 in BMS
   *** times: 0 24+ 48++ hr
   *** reqs: Tp53 [KO]
   *** source: 24076372-Fig-6b
~~~

The **assay** contains at least two parts, the process being measured and the method used to detect it. In datum #2 the protein expression (prot-exp) of Myc is determined by Western Blot (WB) and in datum #3 the phenotype apoptosis is detected by surface expression of Annexin V which is a molecular marker for apoptosis. An assay designation can also contain sites where a modification takes place, a substrate if the activity of the subject is being measured, or a cellular component if a location is being measured. Datum #1 uses a site designation to say that the phosphorylation of S6k1 occurs on Threonine 412 (T412). Datum #4 uses a substrate designation to say that the substrate used to test for Raf1 activity in an in vitro kinase assay (IVKA) is Mek1. Datum #5 uses a location designation to report that Rela relocates to the nucleus 30 min after addition of Tnf to HELA cells.

Datum #4:

~~~
 *** Raf1[Ab]IP IVKA(Mek1)[32P-ATP] is increased irt IL2 (5 min)
   *** cells: CTLL2 in BMS “IL2-deprived”
   *** enhanced by: Wortmannin [pretreatment]
   *** source: 7760801-Fig-9
~~~

Datum #5:

~~~
 *** Rela[Ab] locatedin(nucleus)[IHC] is increased irt Tnf (30 min)
   *** cells: HELA in BMS
   *** inhibited by: SN50 [pretreatment]
   *** source: 10655476-Fig-6c
~~~

When there is a treatment, the **change** can be increased, decreased, unchanged, or detectable-but unchanged. If there is no treatment, detectable or undetectable is used. Datums #1-5 report changes to either the subject or the cellular phenotype. Datum #6 reports that endogenous S6k1 is phosphorylated in T412 in growing cells.

Datum #6:

~~~
 *** S6k1[phosAb] phos(T412)[phosAb] is detectable
   *** cells: MCF7 in BMS
   *** reqs: Mtor [RNAi]
   *** source: 18339839-Fig-2c
~~~

The **stimulus** element has three parts. The treatment is the thing added to or subtracted from the environment. The treatment type is an identifier used to distinguish between adding something directly to the culture supernatant (“irt”, in response to, datums #1-5,8-10), transfecting in a protein (“itpo”, in the presence of, datums #11,12), or deleting a protein by knockout procedures (“itao”, in the absence of, datum #7). If an assay does not use cells, the identifier is “by” to inform us that an interaction is direct and not due to other proteins in the system (datums #13,14). The last treatment element is time. Treatment time is only used for addition of something to the supernatant (type “irt”). Times are not used for “itpo” or “itao” treatments because the time for a protein to be expressed or removed varies with the transfection system used. The time designation refers to the time it takes for a change to occur in the state of protein in a cell. We do not use times in “by” treatments because “by” is only used for reactions that take place in the absence of cells. If a series of times are used, a times line is created. Each time is annotated with zero or more plusses (+) to indicate the change relative to the baseline using + to represent the relative change (datums #3,8).

Datum #7:

~~~
 *** Eif4ebp1[phosAb] phos(T36/T45)[phosAb] is increased itao Tsc2 [KO]
   *** cells: mEFs in BMLS
   *** partially inhibited by: AA-deprivation [addition]
   *** partially inhibited by: AA-deprivation + CHX [addition]
   *** inhibited by: Rapamycin [addition]
   *** source: 15772076-Fig-4d
~~~

**Figure 4:**
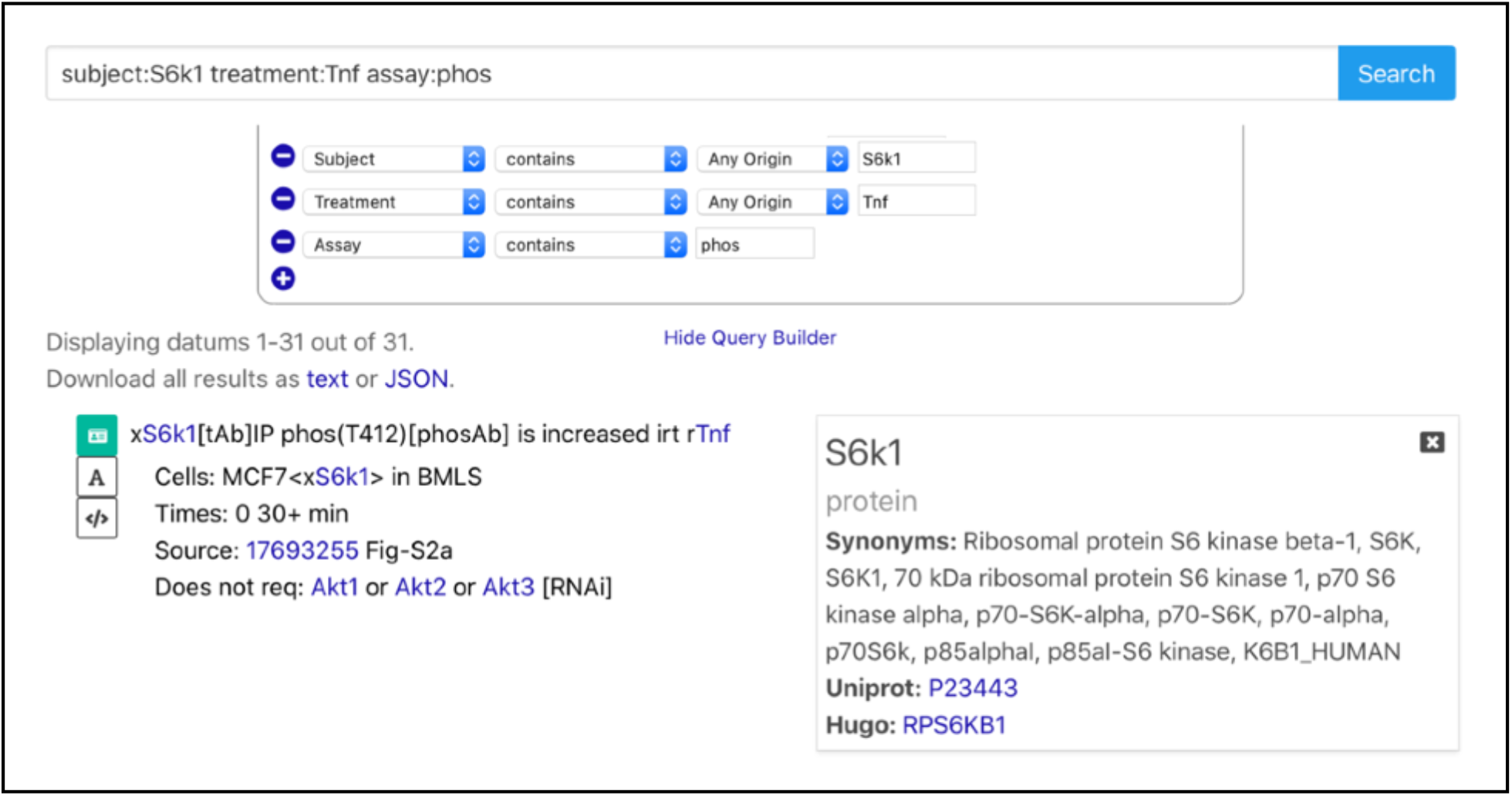
A screenshot of the datum query interface showing one datum in simplified text format and the protein information box for S6k1.

The environment is a description of the starting state of an experiment. It describes the cells used, any exogenously expressed proteins or lack of proteins due to knockouts or knockdowns, and the medium used. Datum #1 is an example showing that the MCF cells have been transfected with a plasmid expressing S6k1. The medium is usually “BMS” (basal medium with serum) to show that a complete growth medium is used, or “BMLS” (basal medium with low serum), which is a catchall term used to represent an attempt to make cells quiescent by removing some or all serum. Currently, free text is used to record any treatments to the cells before the start of the experiment (datums #4, 8, 9) or when a certain subfraction of the cell is used (datum #10).

Datum #8:

~~~
 *** HistH3[phosAb] phos(S11)[phosAb] is decreased irt Nocodazole-removal (times)
   *** cells: HELA in BMS “Synchronized in M by Thymidine-Nocodazole”
   *** times: 0++ 3++ 6 9 12 15+ 18++ 21+ 24 hr
   *** source: 12912980-Fig-1a
~~~

Datum #9:

~~~
 *** Txnip[Ab] copptby[WB] Txn[Ab] is decreased irt Imiquimod (2 hr)
   *** cells: THP1 in BMS “differentiated for 3 hr with PMA”
   *** reversed by: APDC [pretreatment]
   *** source: 20023662-Fig-2b
~~~

Datum #10:

~~~
 *** Hif1a[Ab] oligo-binding[ELISA] is increased irt Hypoxia (6 hr)
   *** cells: NR8383 in BMLS “nuclear-extracts”
   *** source: 14525767-Fig-6a
~~~

Datums can contain additional information in the form of “extras”. Extras report the effects of a system perturbation on the change element. The perturbation can be the removal of a protein by knockout or knockdown, addition of chemical inhibitors, or substitution of a protein with a modified version. The beauty of an extra is that it adds another dimension to the cause and effect reported in the first line. Most curation languages allow you to say that S6k1 is phosphorylated on T412 in response to addition of Tnf. Some have places to say which cells were used and that the cells were harvested after 30 min. But how does one say that it does not happen if Ikk2 is removed from the cells using RNAi (datum#1)? Examples of extras can be seen in datums #1-7,9,11-15,17.

Extras can be used for more than just knockout experiments. Datum #11 says that the phosphorylation of an immunoprecipitate of expressed Mek1 on S218 and S222 in increased on transfection with constitutively active Pak1. This might imply that Pak1 directly phosphorylates Mek1 on S218/S222. But the requirement for Mek1 kinase activity, as seen by the inhibition by substitution of Mek1 with a kinase dead (K97A) version, tells a different story.

Datum #11:

~~~
 *** xMek1[tAb]IP phos(S218/S222)[phosAb] is increased itpo xPak1(T423E)“CA”
   *** cells: rEFs<xMek1> in BMLS “in suspension”
   *** inhibited by: xMek1(S298A) [substitution]
   *** inhibited by: xMek1(K97A)“KD” [substitution]
   *** source: 17314031-Fig-2a
~~~

A datum from the same experiment shows that phosphorylation of Mek1 on S298 does not require Mek1 kinase activity (datum #12).

Datum #12:

~~~
 *** xMek1[tAb]IP phos(S298)[phosAb] is increased itpo xPak1(T423E)“CA”
   *** cells: rEFs<xMek1> in BMLS “in suspension”
   *** inhibited by: xMek1(S298A) [substitution]
   *** unaffected by: xMek1(K97A)“KD” [substitution]
   *** source: 17314031-Fig-2a
~~~

The authors hypothesize that phosphorylation of Mek1 by Pak family kinases induced Mek1 autophosphorylation on the S218/S222 activation loop sites. They tested their theory using an in vitro kinase assay to remove confounding factors such as the upstream Raf family kinases which phosphorylate Mek1 on S218/S222.

Datum #13:

~~~
 *** rMek1[phosAb] phos(S298)[phosAb] is increased by rPak3
   *** cells: none
   *** unaffected by: rMek1(K97A)“KD” [substitution]
   *** inhibited by: rMek1(S298A) [substitution]
   *** source: 17314031-Fig-4a
~~~

Datum #14:

~~~
 *** rMek1[phosAb] phos(S218/S222)[phosAb] is increased by rPak3
   *** cells: none
   *** inhibited by: rMek1(K97A)“KD” [substitution]
   *** inhibited by: rMek1(S298A) [substitution]
   *** source: 17314031-Fig-4b
~~~

As seen in datums #13 and #14 phosphorylation Mek1 on S218/S222 but not S298 requires Mek1 kinase activity, confirming the autophosphorylation theory.

## 4. ENTITIES or THE NAMING OF THINGS

Our model of intracellular signal transduction (http://pl.csl.sri.com/stm8-guide.html) uses a formal system based on rewriting logic called Pathway Logic (http://pl.csl.sri.com/index.html) (1). In this system, each term in the model rules has a unique defined representation. The datums that serve as evidence for the rules use the same defined vocabulary. The terms are given unique names, and not accession numbers, so the user always knows that any terms with the same name represent the same entity in either the model rules or datums. In most cases we use the HGNC symbol with only the first letter in upper case for proteins. Because the focus of the model is on proteins, the name of the gene is the name of the protein followed by “-gene”. This eliminates the confusion caused when the name of a protein is different from its gene symbol. We have also renamed the cell lines using all upper case letters with no punctuation to make them more usable by both humans and computers. For primary cells, a lower case prefix is used to represent the species of origin (h for human, m for mouse, and r for rat, such as mEFs in datum #7, rEFs in datum #11). In some cases cases a term can be mapped to a database. We use UniProt (https://www.uniprot.org) as an anchor for proteins, HGNC (https://www.genenames.org) for genes, Cellosaurus (https://web.expasy.org/cellosaurus/) for cell lines, and Pubmed (https://pubmed.ncbi.nlm.nih.gov) for sources. In cases such as assays, detection methods, and primary cells, we provide our own definitions as specified in the Datum Dictionary at https://pl.csl.sri.com/DatumDictionary/pages/Glossary.html.

Sometimes the exact protein cannot be identified. This happens when an antibody that is used to identify the protein is not specific to one protein. We use the category “family” to indicate a set of proteins that are identified by the same antibody. For example, many antibodies produced to detect Akt1 also see Akt2 and Akt3. When those antibodies are used we use the family name “Akts” to say that we cannot determine which family member(s) is detected. Often a phosphorylation site in a family protein cross reacts with that of another member of the family. Usually we use the amino acid and sequence number to represent a site (e.g., T308 for the protein Akt1). A phosphospecific antibody is produced by immunizing animals with a synthetic phosphopeptide corresponding to residues around the phosphorylated residue. The other family members have identical sequences around that site but the location in the full protein is different (T309 for Akt2 and T307 for Akt3). In these situations we give the site a name using the amino acids in that site, for example:

~~~
Akts[phosAb] phos(KTF)[phosAb] is increased irt Egf (5 min)
~~~

Another common situation is the use of a name for a complex made up of combinations of variable proteins. An example is AMPK which is a trimer of one catalytic alpha chain (Ampka1 or Ampka2), one beta regulatory subunit (Ampkb1 or Ampkb2), and one gamma regulatory subunit (Ampkg1, Ampkg2, or Ampkg3). These are classified as “composites” to indicated that the subject is neither a single protein nor a complex consisting of defined entities.

A protein often has additional attributes such as modifications and mutations and whether it is endogenous or artificially expressed. The name of an expressed protein has the prefix “x” whereas an endogenous protein has no prefix. Mutations in expressed proteins are depicted in parentheses and use the convention *original amino acid/sequence number/new amino acid*. This notation is useful when describing the effect of a mutation on a biological process. For example the datum #15 tells us that the in vitro kinase activity of Ulk1 expressed in HEK293 cells is increased in response to Glucose-deprivation for 4 hours. Mutation of amino acids serine 317 and 777 to alanine abrogates that response.

Datum #15:

~~~
 *** xUlk1[tAb]IP IVKA(Atg13)[32P-ATP] is increased irt Glucose-deprivation (4 hr)
   *** cells: HEK293<xUlk1> in BMS
   *** inhibited by: xUlk1(S317A/S777A) [substitution]
   *** source: 21258367-Fig-3f
~~~

Modifications to proteins are represented by curly brackets. These are used when the subject is detected by an antibody against the modified form of a protein, as seen in datum #16 which says that Rb1 phosphorylated on S612 coprecipitates with E2f1.

Datum #16:

~~~
 *** Rb1{phos(S612)}[phosAb] copptby[WB] E2f1[Ab] is detectable
   *** cells: MOLT4 in BMS
   *** source: 17380128-Fig-2c
~~~

Although the model we are building focuses on human proteins there are occasions when we need to look at protein from other species. This is especially important in the actions of viral proteins in human cells. We use the convention of using *protein!species* for proteins from non-human species. Datum #17 describes the inhibition of Ifnb1 gene expression by the transfection of the Ns protein from Influenza A virus. In addition, it shows that the N-terminal domain (Ns1!IAV(1-80)) is responsible for the inhibition.

Datum #17:

~~~
 *** Ifnb1-gene promo-reporter[CAT] is increased itpo xRigi
   *** cells: A549 in BMS
   *** inhibited by: xNs1!IAV [addition]
   *** inhibited by: xNs1!IAV(1-80) [addition]
   *** unaffected by: xNs1!IAV(81-230) [addition]
   *** source: 17053203-Fig-5a
~~~

## 5. CURATION

As stated earlier, the purpose of collecting datums is to make rules for a model of signal transduction. As anyone who designs experiments knows, there is no way to search PubMed for experiments. We have found that the best way to find experiments is to start at the level of statements about results made in either review articles or the introduction of a data paper. Instead of using the statements directly (the method used by most pathway models), we download the paper referred to in the statement, collect the datums, and, if the data support the statement, we make a rule. Figure 2 shows the process we use to find experiments.

Our curation methods focus on the data presented in the figures and tables. The required information is found in the captions, the results, and the methods (including methods and figures from online supplementary information). For each figure or table, we find as many datum elements as possible, including negative data; all datum elements except extras are required. We use specific terms when an expected element cannot be found in the paper to clarify that the required information was not provided in the paper, as opposed to the curator not recording it. If the site(s) detected by an antibody to a phosphorylated protein is not found in the paper, “sitenr” is used. If the time of a treatment is not supplied, “tnr” is used. If an element is not in the paper the field is marked as “not reported”. Often the authors will not repeat some of the methods used in previous publications and direct the reader to look there. These are also marked “not reported” unless the curator has a reason to search for them.

Our STM model focuses on human cells. Unless the reference indicates the use of a particular isoform or spice variant, the canonical sequence of the human protein in UniProt is used for the sequence numbers in modifications and mutations sites. If mouse proteins or cells are used in experiments, we translate the sites for modifications and mutations into the analogous sites in human proteins so we can compare similar experiments performed in different cells.

The “change” element is the only element that requires an interpretation by the curator. This is not a problem when the data is numerical and statistics are reported; we accept any changes that have a p-value of less than or equal to 0.05. Because of ambiguity regarding “biological replicate” (sometimes it means replicate wells, sometimes independent experiments), we assume that reported statistical significance exists between groups within a single experiment. For this reason, every instance of an experiment is collected in datum form, even if the result has already been demonstrated elsewhere in the paper. The datum format simplifies the detection of contradictions between independent experiments performed in the same paper and the collection of multiple datums reporting similar results to strengthen evidence.

Determining the change becomes problematic when the results are presented in images as is the case for two commonly used detection methods. The first method is immunocytochemistry which uses images to depict changes in the intensity and location of a protein within a cell. The second method is the western blot, in which changes in intensity and the location of a protein on a gel provide information about post-translational modifications. We rely on the expertise of the curator to determine whether a change is significant.

Figure 3 is an example of a western blot from Figure 3f in PMID:17693255 (9) which was used as the source for datum #1. The subject is S6k1 expressed in MCF7 cells and precipitated using an antibody to the tag on the expressed protein (“tAb” is used as a handle). The immunoprecipitate is separated on a SDS-PAGE gel and immunoblotted with an antibody to S6k1 phosphorylated on T412. Here the curator agreed with the authors that the intensity of the phosphorylation was increased in the cells treated with Tnf for 30 minutes although the amount of total S6k1 proteins was unchanged (compare lane 1 with lane 2). There is no increase in the absence of Ikk2 (compare lane 6 with lane 7), so the extra saying that Ikk2 is required is justified.

## 6. DATUM QUERY INTERFACE

The best way to access, filter, and download datums is to use the Datum Query Interface (DQI, http://datum.csl.sri.com). The search engine can be used by either typing a query directly into the search box or by using the query builder which can be accessed by clicking on “Show Query Builder” under the search box. The results are displayed in a simplified text format showing the first line, the environment line, a line containing the treatment times if applicable, the source line, and any extras comments. Any item colored blue can be clicked on for more information. Clicking on a protein will bring up a box containing the Uniprot ID with a link to the Uniprot protein record, the HGNC symbol with a link to the HGNC record, and synonyms. Clicking on a chemical will bring up a box containing synonyms, a putative activity, and a link to the PubChem record. Clicking on the PubMed ID in the source field opens the PubMed record.

Figure 4 contains a screenshot of the search results using the DQI. The datum shown was one of 31 found by asking the interface for a datum whose subject contains the protein S6k1 (either endogenous or exogenous), the treatment Tnf, and the assay “phos”. The box on the right is the result of clicking on xS6k1 showing synonyms and providing links to the PubMed and HGNC records.

Typing a term into the search box will find all datums that contain it. The search can be limited to specific fields within the datum by prefixing the term with tags representing those fields or expanded by using a wild-card symbol (*). The query builder can be used to provide the tags, a set of conditions, and a choice of terms, if they are limited. A link (http://pl.csl.sri.com/datumkb.html) is provided to an explanation of how to read a datum, a set of example queries, and a link to the Datum Dictionary, which contains the definitions of assays, detection methods, and other datum elements.

## 7. USES FOR DATUMS

Datums are more than just evidence for our Pathway Logic model; they are often useful for finding details when designing experiments. It helps to know in advance which cell lines will respond to a stimulus or which proteins are active when overexpressed. Datums can find similar experiments to the current design in process and/or show the results obtained using a different cell line. Because datums are always mapped to experiments (and not assertions), you can go to the source and extract finer details such as lysis buffer components for protein extraction and fixatives used for immunohistochemistry.

Datums were used by coworkers who were developing ways to identify potential threats from protein sequences. One of their assays was to express the proteins in cultured cells and measure the effect on survival. When they needed to know the control proteins that impair the growth of cells when overexpressed, a search in the DQI identified them. A common phenotype of cell death is an increase in apoptosis, but the researchers were only interested in the effects of expressed proteins; the DQI allowed limiting the search to the treatment type “itpo”. The query

~~~
assay:apoptosis change:increase treattype:itpo
~~~

produced 80 datums. From these search results, the researchers extracted 22 normal proteins, 12 mutated proteins, and 3 protein combinations that could serve as controls.

In another project studying potential host-virus interactions, we collected experiments where viral proteins were shown to interfere with signal transduction. The query: “viral:*” was used to find all datums involving viral proteins. The search was limited to protein interactions by using “assay:copptby”. The result included datums describing experiments in which human proteins coprecipitated with viral proteins or viral proteins interfered with interactions between human proteins.

Datums can be helpful for determining the reproducibility of a result. The query:

~~~
subject:Erks assay:phos treatment:“Egf”
~~~

produces 372 datums in which the phosphorylation was marked “increased”, 9 as “detectable but unchanged”, and 1 as “unchanged”. Examining the cell lines used in the contradictory datums revealed 7 that contained activating mutations of the Braf protein and 2 that contained activating mutations of the Kras protein. Either of those mutations would cause Erks to be constitutively phosphorylated and to have a high basal background. The last datum looked at nuclear extracts. No total Erk protein controls were performed, so it is unclear whether there was enough protein in the extracts to detect phosphorylation. In contrast, the same query using P38s or Jnks as subjects showed much less reproducibility (5 of 24 unchanged for P38s, and 6 of 20 unchanged for Jnks). These results increase our confidence that Erks are phosphorylated in response to Egf treatment but less so for P38s or Jnks.

## 8. CONCLUSION

Datums were originally designed to be used as evidence for rules in a Pathway Logic model of intracellular signal transduction (STM8, http://pl.csl.sri.com/online.html) (1,10). They have since been used as a training set and target information structure in developing natural language processing tools for reading biological papers (11–13). Burns et al. (11) used datums to provide the annotation scheme needed to to automate the process of extracting experimental observations from articles based on subdividing the text of articles into relatively compact discourse elements pertaining to a single experiment. Freitag and Niekrasz (12) used datums to evaluate their method of extracting biological information using composite frames constituted from multiple sentences. Freitag and colleagues (13) used the various datum elements described in the section on Datum structure to construct reference frames from experimental information distributed across a document.

Convention specifies that an experimental result must be published in a journal indexed in PubMed before it is considered valid, but no tools are currently available to search for experiments. Until there are, we will continue to curate and store them manually. We have used datums often to answer these questions: I have a hypothesis. I could test it by doing this experiment. Has anyone ever done it or something like it? What cells/reagents/time range/ concentration range should I use? Although the DatumKB does not cover all the publications about intracellular signal transduction in PubMed, it contains almost 100,000 datums from 3314 papers. We believe that the contents of the DatumKB will be helpful to bench scientists today and the development of new methods to automate the extraction of biomolecular information from biological research literature in the future.

## 9. FUNDING

This work was supported by the National Institutes of Health [CA112970, GM068146], the Defense Advanced Projects Agency [W911NF-14-C-0108, W911NF-14-2-0020], Intelligence Advanced Research Projects Activity [W911NF-17-C-0051], and the National Science Foundation [2034772, IIS-0513857].

## Notes

https://datum.csl.sri.com

http://pl.csl.sri.com/online.html

### Competing Interest Statement

The authors have declared no competing interest.

